# Site-specific cholesterol depletion therapy for gastric cancer

**DOI:** 10.64898/2026.07.15.738777

**Authors:** Simona Kavrakova, Ashutosh Sharma, Elena Ristevska, Xingyu Guo, Julian Zalejski, Wonhwa Cho

**Author notes:** These authors equally contributed to this work (Simona Kavrakova, Ashutosh Sharma). Corresponding author: Wonhwa Cho, Department of Chemistry, University of Illinois Chicago, Chicago, IL 60607, USA;. Lead contact: Wonhwa Cho, Department of Chemistry, University of Illinois Chicago, Chicago, IL 60607, USA.

## Abstract

Altered cholesterol metabolism is a recognized hallmark of cancer, but systemic modulation has not yet delivered significant clinical results. Accumulating evidence shows that cholesterol plays distinct roles across diverse cellular membranes, suggesting that site-specific modulation may produce superior therapeutic outcomes. Cholesterol is associated with gastric cancer (GC), but the mechanistic link is complex and no effective cholesterol-targeted therapy has been developed. Here, we report that cholesterol levels in GC cells are site-specifically elevated in the inner leaflet of the plasma membrane (IPM). This elevated IPM cholesterol constitutively activates Wnt-β-catenin signaling to drive cell survival and proliferation. Mechanistically, Niemann-Pick C1-like 1 (NPC1L1), which is highly expressed in GC patient tissues and cell lines, acts as a cholesterol flippase to raise IPM cholesterol levels, facilitating ligand-independent β-catenin signalosome formation. Ezetimibe, a clinically approved NPC1L1 inhibitor, blocks this flippase activity, lowers IPM cholesterol levels, and suppresses β-catenin signaling. Ezetimibe treatment induces apoptosis in GC cells while sparing normal primary gastric epithelial cells, which exhibit low levels of NPC1L1 and IPM cholesterol. Collectively, these results suggest that site-specific modulation of cellular cholesterol is a viable approach to developing safe and effective therapies for cancers linked to local cholesterol elevation.

## INTRODUCTION

Lipids are essential for cell survival as they serve as an energy source and cellular building blocks (1, 2). Lipids also control numerous cellular processes by modulating structure, function, localization, and mutual interaction of cellular proteins (3, 4). To drive cell growth, proliferation and survival, cancer cells upregulate diverse lipids through reprogramming of lipid metabolism and transport (1, 2). Of all lipids, cholesterol is most extensively linked to cancer (5, 6). Normally, cellular cholesterol homeostasis is tightly regulated by a complex mechanism that controls its import, intracellular transport, storage, synthesis, metabolism, and export (7). In tumor cells, many proteins involved in cholesterol biosynthesis, import, and storage are upregulated, as cholesterol is essential for cancer cell survival (8). Consequently, inhibitors of cholesterol metabolism—most notably statins—have been extensively evaluated for their potential anti-tumor activities (9); however, they have not produced consistent clinical benefits despite their ability to lower the systemic cholesterol levels (10). These results underscore the need for thorough investigation of the cell type-specific and site-specific roles of cholesterol in the tumor microenvironment with an emphasis on the spatiotemporal molecular mechanisms directly linking cholesterol to cancer cell progression. They also call for a new therapeutic strategy that targets site-specific cholesterol function and levels, rather than general cholesterol metabolism and homeostasis.

It has long been known that the large majority of cellular free (or unesterified) cholesterol is located in the plasma membrane (PM) (11, 12). Recent advances in mass spectrometry-based lipidomics and biosensor-based quantitative microscopic imaging have revealed that cholesterol exists in spatially and functionally distinct pools across subcellular membranes beyond PM, including lysosomes, mitochondria, and the endoplasmic reticulum (13). A large body of literature describes the involvement of cholesterol in cancer signaling via cholesterol-rich lipid rafts within the PM (14, 15). However, the existence and functional significance of lipid rafts in living cells remain controversial, and the specific and clinically effective modulation of lipid rafts has not yet been reported. Recent studies have suggested an alternative mechanism for the site-specific signaling function of cholesterol in the PM. Notably, simultaneous quantitative ratiometric imaging of cholesterol in the two leaflets of PM (16, 17) showed that the cholesterol level in the inner leaflet of PM (IPM) ([Chol]*_ipm_*) is kept much lower (i.e., <0.5 mole%) than in the outer leaflet (OPM) (i.e., >30 mole%) in normal epithelial cells (16, 18). Similar transbilayer cholesterol asymmetry in the PM was found in HEK293 GnTI cells (19) and erythrocytes (20) by independent methods. In cancer cells, [Chol]*_ipm_* is site-specifically elevated in a stimulus-independent manner (18), presumably through re-wiring of cholesterol homeostasis (21). This local cholesterol enrichment constitutively activates a subset of signaling proteins via unique and specific cholesterol-protein interaction, thus driving cancer type-specific pro-tumorigenic signaling networks (18). For example, elevated [Chol]*_ipm_* in *APC*-truncated colorectal cancer (CRC) cells directly binds a scaffolding protein, dishevelled (Dvl), and drives ligand-independent Wnt–β-catenin signaling (16, 22). Since these signaling networks are absent in normal cells with low [Chol]*_ipm_*, they present attractive targets for cancer-specific drug development (18). Site-specific signaling function of lysosomal cholesterol has been also reported (23).

Since cholesterol is indispensable for cell membrane integrity and plays pleiotropic roles in physiological processes as a signaling molecule (13), systemic depletion of cellular cholesterol would exert deleterious effects on normal cells. For example, common cholesterol-lowering agents such as statins and m-β-cyclodextrin severely damage the morphology and physiology of normal cells at the doses necessary for suppressing cancer cells (24). To date, no pharmacological inhibitor has been reported to site-specifically deplete cellular cholesterol. This work aimed to develop a site-specific cholesterol depletion therapy with a focus on selectively lowering elevated IPM cholesterol in cancer cells. We focused on gastric cancer (GC) as a model in this study because we found that GC cells exhibited site-specific, high-level accumulation of cholesterol within the IPM. Elevated [Chol]*_ipm_* in GC cells constitutively activated Wnt-β-catenin signaling to drive cell survival and proliferation. We also found that Niemann-Pick C1-like 1 (NPC1L1), which was highly expressed in all tested GC cells, acted as a cholesterol flippase to raise the IPM cholesterol levels and that ezetimibe, a clinically approved inhibitor of NPC1L1, blocked its flippase activity and consequently site-specifically lowered the [Chol]*_ipm_* and suppressed the β-catenin signaling in GC cells. Ezetimibe potently induced apoptotic death of GC cells, while sparing normal primary gastric epithelial cells that exhibited low levels of NPC1L1 and IPM cholesterol.

## MATERIALS AND METHODS

### Materials

Ezetimibe, U18666A, Avasimibe, cholesterol, and other small-molecule inhibitors were purchased from Sigma-Aldrich unless otherwise indicated. Stock solutions of ezetimibe, avasimibe, and U18666A were prepared in dimethyl sulfoxide (DMSO) and stored at −20 °C. Working concentrations were freshly prepared by dilution in cell culture medium, with the final DMSO concentration not exceeding 0.1% (v/v). Phospholipids used for membrane and vesicle-related assays were obtained from Avanti Polar Lipids. All tissue culture media, fetal bovine serum (FBS), antibiotics, and supplements were purchased from Thermo Fisher Scientific. Antibodies against NPC1L1, non-phosphorylated (active) β-catenin (Ser33/37/Thr41), cleaved caspase-3, and GAPDH were obtained from Cell Signaling Technology. Horseradish peroxidase (HRP)-conjugated secondary antibodies were purchased from Jackson ImmunoResearch. Fluorescent cholesterol sensors used for subcellular cholesterol quantification were synthesized and validated as described previously, with targeting motifs enabling selective reporting of cholesterol in defined membrane compartments. All chemicals were of analytical grade or higher.

### Cell Culture, Cell Transfection, and siRNA Knockdown

Human CRC cell lines (HCT15, Caco-2) and GC cell lines (NCI-N87, AGS, KATOIII, Hs746T) were obtained from the American Type Culture Collection (ATCC). Human gastric epithelial primary cells (HGaEpC) were purchased from Cell Applications Inc. and used as non-malignant controls. Cell line identities were confirmed by short tandem repeat (STR) profiling, and all cells were routinely tested and confirmed negative for *Mycoplasma* contamination. CRC cell lines were cultured in Dulbecco’s modified Eagle’s medium (DMEM) supplemented with 10% (v/v) FBS and 1% penicillin–streptomycin. GC cell lines were cultured in RPMI-1640 supplemented with 10% FBS. HGaEpC cells were cultured in GI Epithelial Cell Defined Culture Medium (Cell Applications Inc.) according to the supplier’s recommendations. Cells were maintained at 37 °C in a humidified incubator containing 5% CO₂ and used for experiments between passages 5 and 20.

For transient transfection, cells were seeded at 50–60% confluency and transfected using Lipofectamine 3000 according to the manufacturer’s protocol. For single-molecule imaging experiments, plasmids encoding fluorescently tagged Frizzled or Axin were transfected at low DNA concentrations to ensure sparse expression suitable for single-particle tracking. For siRNA-mediated knockdown, cells were transfected with validated siRNA duplexes targeting *NPC1L1* or non-targeting control siRNA (Integrated DNA Technologies) at a final concentration of 20–40 nM using Lipofectamine RNAiMAX. Cells were harvested 48–72 h post-transfection, and knockdown efficiency was confirmed by immunoblot analysis.

### Quantitative cholesterol imaging

Spatiotemporally resolved *in situ* quantification of intracellular cholesterol in mammalian cells was performed using ratiometric cholesterol sensors, WCR-YDA (all intracellular cholesterol) and DAN-D434A (cholesterol in the OPM), as described previously (16, 25). Briefly, WCR-YDA and DAN-D434A were prepared by chemical conjugation of the YDA (Y415A /D434A/A463W) and D434A mutants of the D4 domain of perfringolysin O with solvatochromic fluorophores, WCR (26) and acrylodan (DAN; MedChemExpress), respectively, and calibrated using giant unilamellar vesicles containing POPC/POPS/cholesterol (80-*x*:20:*x; x* = 0-40 mole%). WCR-YDA was microinjected into cells, and the cholesterol concentration in the IPM and lysosomes was determined using in-house programs written in MATLAB. DAN-D434A was added to the cultured medium for cell surface cholesterol quantification. To validate the quantification results from WCR-YDA, intracellular cholesterol quantification was also performed with an orthogonal cholesterol sensor, WCR-*e*Osh4 as reported previously (25). The three-dimensional display of local lipid concentration profile was calculated using the Surf function in MATLAB.

### Site-specific cholesterol depletion

Site-specific cholesterol depletion was performed as described with a minor modification (27). For transient expression of Lyn-FRB and FKBP_2_-mTagBFP-WVR for PM targeting, Lyn-FRB and FKBP_2_-mTagBFP-WVR were separately subcloned into the pcDNA3 vector. Equal amounts of the resulting plasmids were transfected into CRC cells using the JetPRIME system (Polyplus-transfection) according to the manufacturer’s protocol. The growth medium was switched to Invitrogen^TM^ Live Cell Imaging Solution next day and the PM targeting of FKBP_2_-mTagBFP-WVR was induced by adding rapalog to the solution to a final concentration of 50 nM. Lysosomal and endoplasmic reticulum (ER) targeting of FKBP_2_-mTagBFP-WVR was performed by the same protocol except that Lamp1-iRFP713-FRB and calnexin-iRFP713-FRB, respectively, were used in lieu of Lyn-FRB.

### Single-Molecule Tracking Analysis

Single-molecule imaging was performed using a custom-built total internal reflection fluorescence microscope (TIRFM) as described previously (18, 22). Human GC cells were plated on the 8-well chambered cover glass (Lab-Tek, Thermo Fisher Scientific) at the density of 1 × 10^5^ for 24 h and Dvl2-EGFP plus Halo-tagged axin1 were co-transfected into cells using the jet-PRIME system (Polyplus-transfection) according to the manufacturer’s protocols. Cells containing mobile protein signals with distinguishable blinking patterns (small dots a few pixels in size) but lacking protein aggregates (i.e., large immobile bodies of fluorescence signal) were selected for analysis. For co-localization analysis, cells showing comparable fluorescence signals in the green (i.e., EGFP) and red (i.e., TMR) channels after labeling were selected for analysis.

For colocalization analysis, dual-transfected GC cells were subsequently labeled with HaloTag® TMR (Promega) the next day and washed extensively according to the manufacturer’s protocols. Two protein molecules were simultaneously tracked and analyzed as described using in-house programs written in MATLAB (22). The images were spatially corrected following the algorithm described previously (28). Co-localization analysis of two molecules was performed with a fixed threshold criterion (i.e., <400 nm) for co-localization (28). The percentage Dvl2 molecules colocalized with FZD7 (or axin1, LRP6) on the PM of CRC cells was calculated from the total colocalization events occurred and displayed as a histogram. Data were fit into a single exponential decay equation (i.e., *P* = *P*_o_ *e^-kt^*) to determine the dissociation rate constant (*k*) values for the protein complexes by non-linear least-squares analysis and the half-life values of co-localization were calculated as *ln*2/*k*.

### Cell Viability Assay

Cell viability was determined using the CellTiter-Glo® 2.0 Luminescent Cell Viability Assay (Promega) according to the manufacturer’s instructions. Briefly, cells were seeded in white, opaque 96-well tissue culture plates at a density of 5 × 10³ cells per well and allowed to attach overnight. Cells were then treated with the indicated concentrations of ezetimibe or vehicle control (0.1% DMSO) for 72 h under standard culture conditions. Cell morphology was routinely monitored by phase-contrast microscopy throughout the treatment period. Following treatment, an equal volume of CellTiter-Glo® 2.0 reagent was added directly to each well. Plates were mixed on an orbital shaker for 2 min to induce complete cell lysis and incubated at room temperature for 10 min to stabilize the luminescent signal. Luminescence was measured using a BioTek microplate reader. Cell viability was normalized to vehicle-treated controls and expressed as the percentage of viable cells relative to control. Dose-response curves and IC₅₀ values were determined by nonlinear regression analysis. Each experimental condition was analyzed in triplicate technical replicates, and all experiments were independently repeated at least three times.

### Western Blot Analysis

Cells were washed twice with ice-cold phosphate-buffered saline (PBS) and lysed in RIPA buffer (Thermo Fisher Scientific) supplemented with protease and phosphatase inhibitor cocktails (Thermo Fisher Scientific). Cell lysates were clarified by centrifugation at 14,000 × g for 15 min at 4°C, and protein concentrations were determined using the Pierce™ BCA Protein Assay Kit (Thermo Fisher Scientific). Equal amounts of protein (20 μg) were resolved by 10% SDS-PAGE and transferred onto 0.45-μm nitrocellulose membranes using a wet-transfer system. Membranes were blocked with 5% bovine serum albumin (BSA) in Tris-buffered saline containing 0.1% Tween-20 (TBST) for 1 h at room temperature, followed by incubation overnight at 4°C with primary antibodies against NPC1L1 (Cell Signaling Technology, Cat. #5058, 1:1000), non-phosphorylated (active) β-catenin (Cell Signaling Technology, Cat. #8814, 1:1000), cleaved caspase-3 (Asp175) (Cell Signaling Technology, Cat. #9661, 1:1000), and GAPDH (Cell Signaling Technology, Cat. #2118, 1:1000). After three washes with TBST, membranes were incubated with secondary antibodies for 1 h at room temperature. Immunoreactive bands were detected using Clarity™ Western ECL Substrate (Bio-Rad Laboratories) and visualized using an Azure imaging system. Band intensities were quantified using ImageJ software, normalized to the corresponding GAPDH loading controls, and expressed relative to the vehicle-treated controls. All immunoblotting experiments were performed using at least three independent biological replicates.

### Colony formation assay

6-well plates were prepared with 3 mL of 0.6% 2-hydroxyethyl agarose (Catalog #39346-81-1, Sigma Aldrich) in the same media as the cells were cultured in, the plates were incubated at 4 C° for 1 hour and transferred to 37 C° for at least 30 min before plating cells. Cells were plated in 1 mL of 0.3% agarose per well in triplicate with 40,000 cells per well with inhibitor. Colonies were allowed to grow until visible, about 3 weeks for MDA-IBC3. Plates were imaged and analyzed with the Nexcelom Celigo Image Cytometer.

### Gene Expression Analysis

The web-based tool Gene Expression Profiling Interactive Analysis v2 (GEPIA2) was accessed to compare the relative mRNA expression levels of *NPC1L1* in stomach adenocarcinoma (STAD) and other cancer tissues against normal tissue samples acquired from the TCGA and GTEx databases. Differential analysis of expressed genes between TCGA tumor samples and paired normal samples from the TCGA and GTEx projects was performed using the limma method. Disease-free survival (DFS) analysis was performed based on gene expression using the log-rank (Mantel–Cox) test. R packages were used for generating all diagrams and statistical analyses.

### RNA Sequencing Analysis

AGS cell samples were submitted to Plasmidsaurus (Berkeley, CA, USA) for RNA sequencing. Plasmidsaurus performed RNA extraction, library preparation, high-throughput sequencing, quality control, read alignment to the reference genome, transcript quantification, and differential gene expression analysis. Differentially expressed genes (DEGs) were reported as log_2_ fold change and false discovery rate (FDR)-adjusted *p* values. A volcano plot was generated in R using the ‘ggplot2’ and ‘ggrepel’ packages. Genes were plotted according to log_2_ fold change and −log_10_(FDR). Genes meeting the thresholds of ∣log_2_ fold change∣ ≥1 and FDR <0.05 were considered significantly differentially expressed. Gene Set Enrichment Analysis (GSEA) was performed using the Broad Institute GSEA software with the MSigDB Hallmark gene set collection. Genes were ranked according to differential expression, and pathway enrichment was evaluated using the normalized enrichment score (NES), nominal *p* value, and FDR *q* value. The top positively and negatively enriched Hallmark pathways, ranked by NES, were visualized as a horizontal bar plot using ‘ggplot2’ in R. Representative enrichment plots for specific pathways of interest were exported using GSEA software. A hierarchically clustered heatmap was generated using Morpheus (Broad Institute). Target genes were selected from the Hallmark Wnt/β-catenin signaling gene set, and normalized expression values (counts per million, CPM) were converted to row-wise Z-scores prior to hierarchical clustering and visualization.

### Statistical Analysis

All data are presented as mean ± standard deviation (SD) unless otherwise indicated. Statistical analyses were performed using Kaleidagraph (Synergy Software). Comparisons between two groups were performed using unpaired, two-tailed Student’s *t*-tests. For multiple group comparisons, one-way ANOVA followed by appropriate post-hoc tests was used. Differences were considered statistically significant at *p* < 0.05.

## RESULTS

### IPM cholesterol levels are site-specifically elevated in gastric cancer cells

Cholesterol has been extensively associated with GC; however, the nature of that association is complex and context-dependent (29–31). To understand the GC cell-specific function of cholesterol, we first determined its subcellular distribution. Filipin has been widely used to estimate the subcellular distribution of cholesterol but this method has major limitations that hamper cholesterol quantification in live cells (32). Furthermore, while many genetically encoded cholesterol probes have been reported (32–34) none allows for robust spatiotemporally resolved quantification of cholesterol. We therefore quantified the local cholesterol concentrations in GC and other control cells by our recently developed quantitative cholesterol imaging technique (16, 25). Spatiotemporally resolved quantification of intracellular cholesterol was performed by ratiometric imaging analysis using two orthogonal cholesterol-specific sensors with high cholesterol affinity, WCR-YDA or WCR-eOsh4 (16, 25), whereas that in the OPM with a lower affinity cholesterol sensor, DAN-D434A (16) (**Fig 1a**). These ratiometric sensors allow for robust spatiotemporally resolved *in situ* quantification of cholesterol in live cells (16, 25)

**Fig 1.**
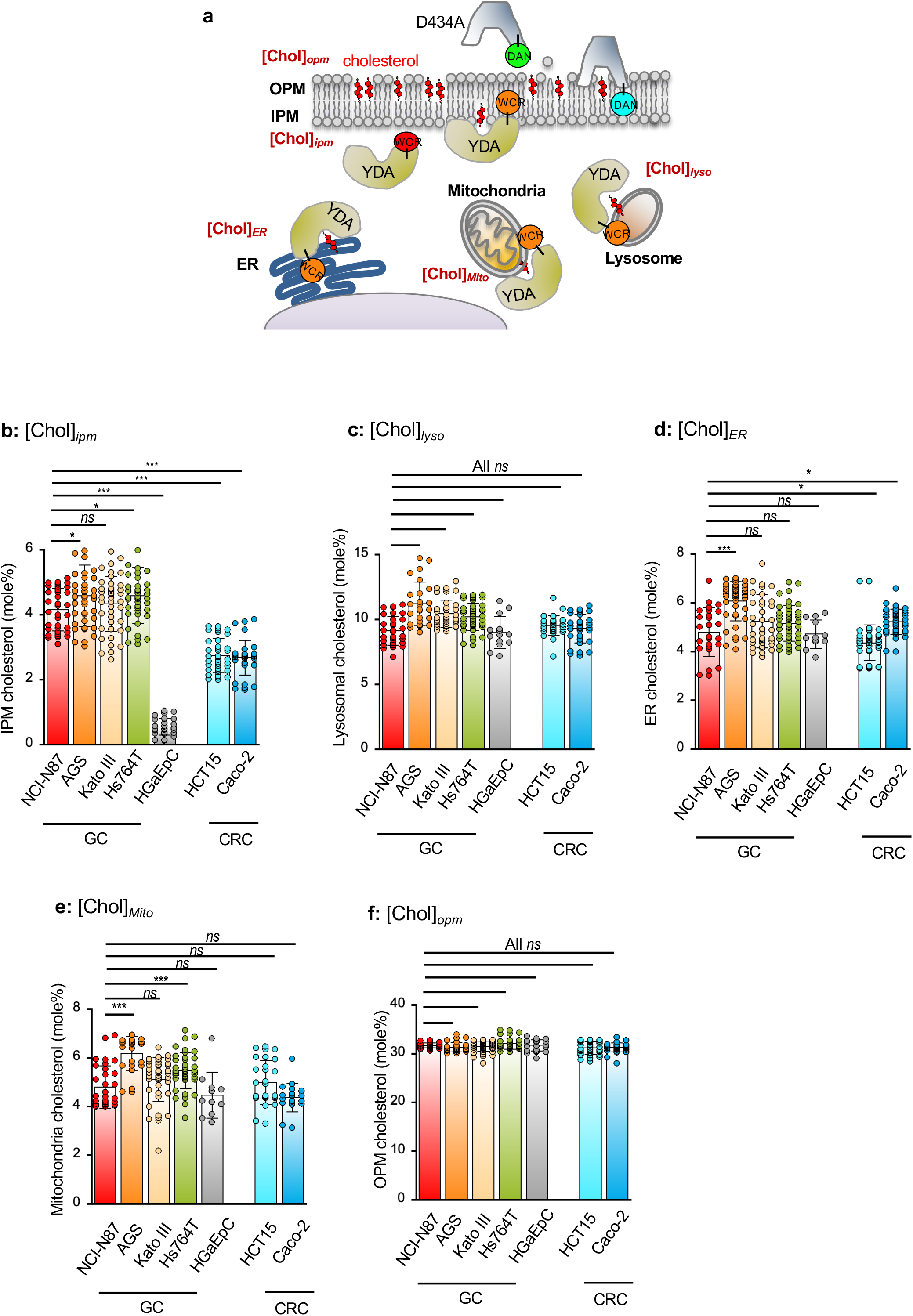
Spatially resolved cholesterol quantification in gastric cancer cells, normal gastric epithelial cells, and colorectal cancer cells. **a**. A general scheme of spatiotemporal cholesterol quantification. The cholesterol concentrations in the cytofacial leaflets of the plasma membranes and organelle membranes were determined using a microinjected ratiometric cholesterol sensor, WCR-YDA, which undergoes a hypsochromic shift (or blue shift) in fluorescence emission upon binding to membranes. The cholesterol concentrations in the outer leaflets of the plasma membrane were determined with another ratiometric sensor, DAN-D434A, which was added to the media. WCR-YDA and DAN-D434A have different cholesterol response ranges and spectral ranges. **b-f**. Spatially averaged cholesterol concentrations in the inner plasma membrane ([Chol]*_ipm_*) (b), lysosomes ([Chol]*_lyso_*) (**c**), endoplasmic reticulum (ER) ([Chol]*_ER_*) (**d**), mitochondria ([Chol]*_Mito_*) (**e**), and the outer plasma membrane ([Chol]*_opm_*) (f). Quantification was performed in triplicate with different samples and *n* ≥ 6 cells per experiment. Values are means ± SD. Statistical analysis was performed with unpaired Student’s t-test. *ns*, *p* > 0.05; *, *p* < 0.05; **, *p* < 0.01; ***, *p* < 0.001.

We previously reported that [Chol]*_ipm_* is kept low (i.e., <0.5 mol%) in primary epithelial cells to restrict their proliferative activity (16, 18). Human gastric epithelial primary cells (HGaEpC) similarly exhibited a low [Chol]*_ipm_* (0.5 ± 0.2 mol%) value. Notably, all tested GC cell lines (NCI-N87, AGS, Kato III, and Hs764T) possessed uniformly and markedly higher [Chol]*_ipm_* values (i.e., >4.0 mol%) than HGaEpC cells (**Fig. 1b**). These values exceeded those found in *Apc*-truncated colorectal cancer (CRC) cells (**Fig. 1b**), a gastrointestinal malignancy where elevated IPM cholesterol characteristically drives constitutive Wnt-β-catenin signaling (18). This local elevation of cholesterol was unique to the IPM, as cholesterol levels in other cytofacial membranes— namely lysosomes ([Chol]*_lyso_*) **(Fig. 1c**), the endoplasmic reticulum (ER) ([Chol]*_ER_*) (**Fig. 1d**), and mitochondria ([Chol]*_mito_*) (**Fig. 1e**)—showed no significant differences among the GC, normal gastric, and CRC cell lines. Similarly, no significant difference was found in the OPM cholesterol levels ([Chol]*_opm_*) among these cells (**Fig. 1f**).

### Elevated IPM cholesterol levels control the survival and proliferation of GC cells

The uniquely elevated IPM cholesterol across GC cell lines suggests that it may play a critical functional role in these cells. To assess the functional importance of IPM cholesterol, we first measured how site-specific depletion of cholesterol affects GC cell viability. The site-specific depletion of cholesterol was achieved by targeting of an engineered cholesterol oxidase (CholOx) to specific membranes via chemically induced protein heterodimerization (27) (**Fig. 2a**), which was subsequently confirmed by ratiometric cholesterol quantification (**Fig. 2b**). Cell viability was then assessed using the CellTiter-Glo **®** luminescent assay. When IPM cholesterol was specifically depleted in the four GC cell lines, cell viability was reduced by more than twofold; however, site-specific depletion of lysosomal and ER cholesterol exerted little to no effect (**Fig. 2c**). These results show that elevated IPM cholesterol plays a major role in GC cancer cell survival and proliferation.

**Fig 2.**
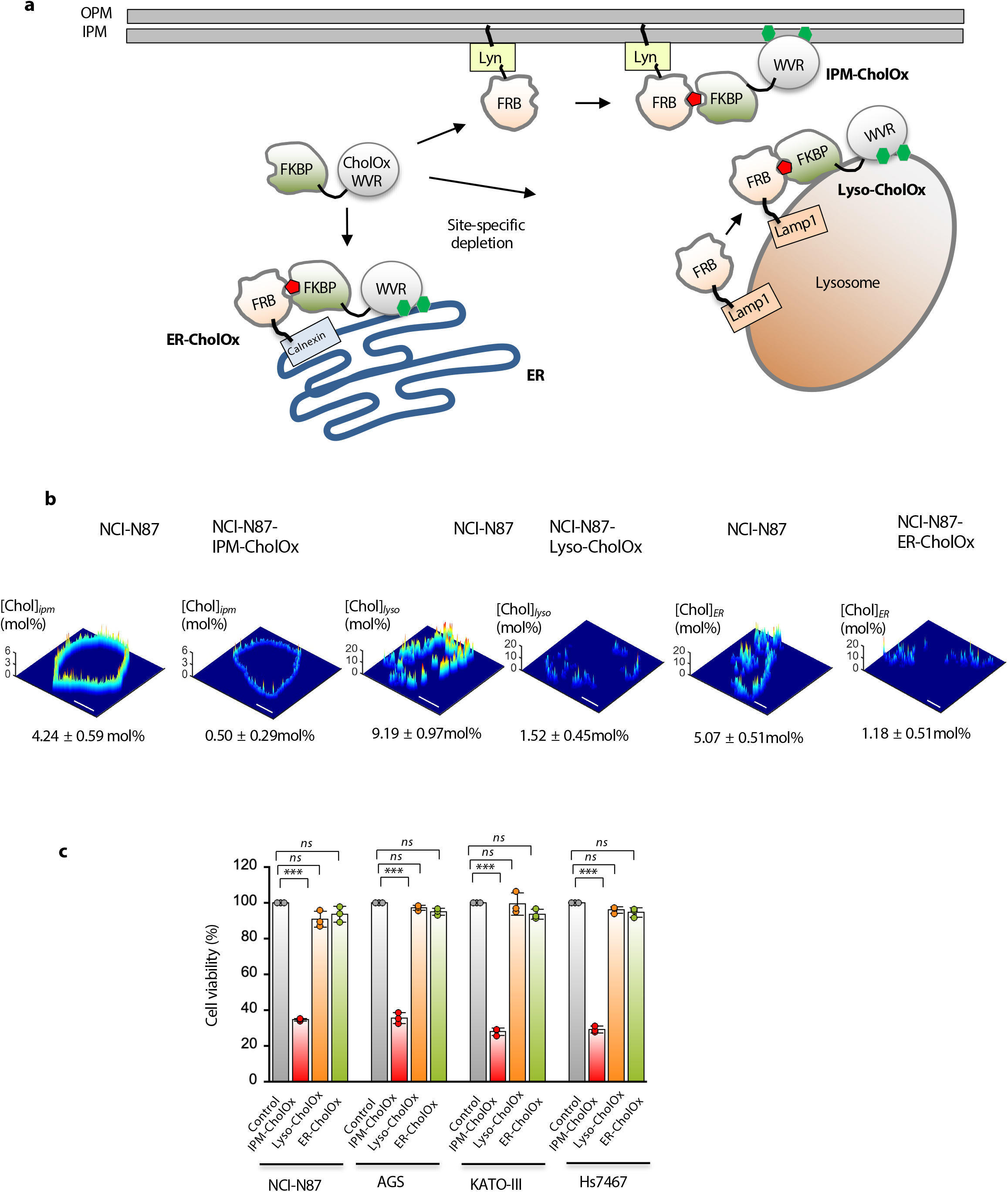
Effects of site-specific depletion of cholesterol on GC cell viability. **(a)** The site-specific cholesterol depletion strategy. The engineered cholesterol oxidase (CholOx-WVR) is site-specifically targeted via rapalog (red)-mediated dimerization of FKBP and FRB. **(b)** Spatiotemporally resolved [Chol]*_ipm_*, [Chol]*_ER_*, and [Chol]*_lyso_* profiles in NCI-N87 cells before and after site-specific cholesterol depletion. The *z*-axis scale indicates the cholesterol concentration (mol%). A spatially averaged concentration (average ± SD from triplicate determinations with >5 cells per measurement) value is shown for each cell type. A pseudo-coloring scheme with red and blue representing the highest and the lowest concentration, respectively, is used to illustrate the spatial cholesterol heterogeneity. Scale bars indicate 10 μm. **(c)** Effects of site-specific cholesterol depletion on GC cell viability. *N* = 3. All values are means ± SD. Statistical analysis was performed with unpaired Student’s t-test. *ns*, *p* > 0.05; *, *p* < 0.05; **, *p* < 0.01; ***, *p* < 0.001.

### Elevated IPM cholesterol levels drive constitutively active Wnt-β-catenin signaling

To investigate the molecular mechanisms by which IPM cholesterol controls GC cell viability, we performed global RNA-sequencing on AGS cells before and after CholOx-mediated cholesterol depletion. Differential expression analysis revealed that four key Wnt/β-catenin target genes were downregulated upon cholesterol depletion (**Fig. 3a**, **3b**). Unbiased Gene Set Enrichment Analysis (GSEA) further identified Wnt/β-catenin as the most significantly suppressed signaling pathway in cholesterol-depleted cells (**Fig. 3c**), driven by widespread downregulation of its pathway components (**Fig. 3d**).

**Fig 3.**
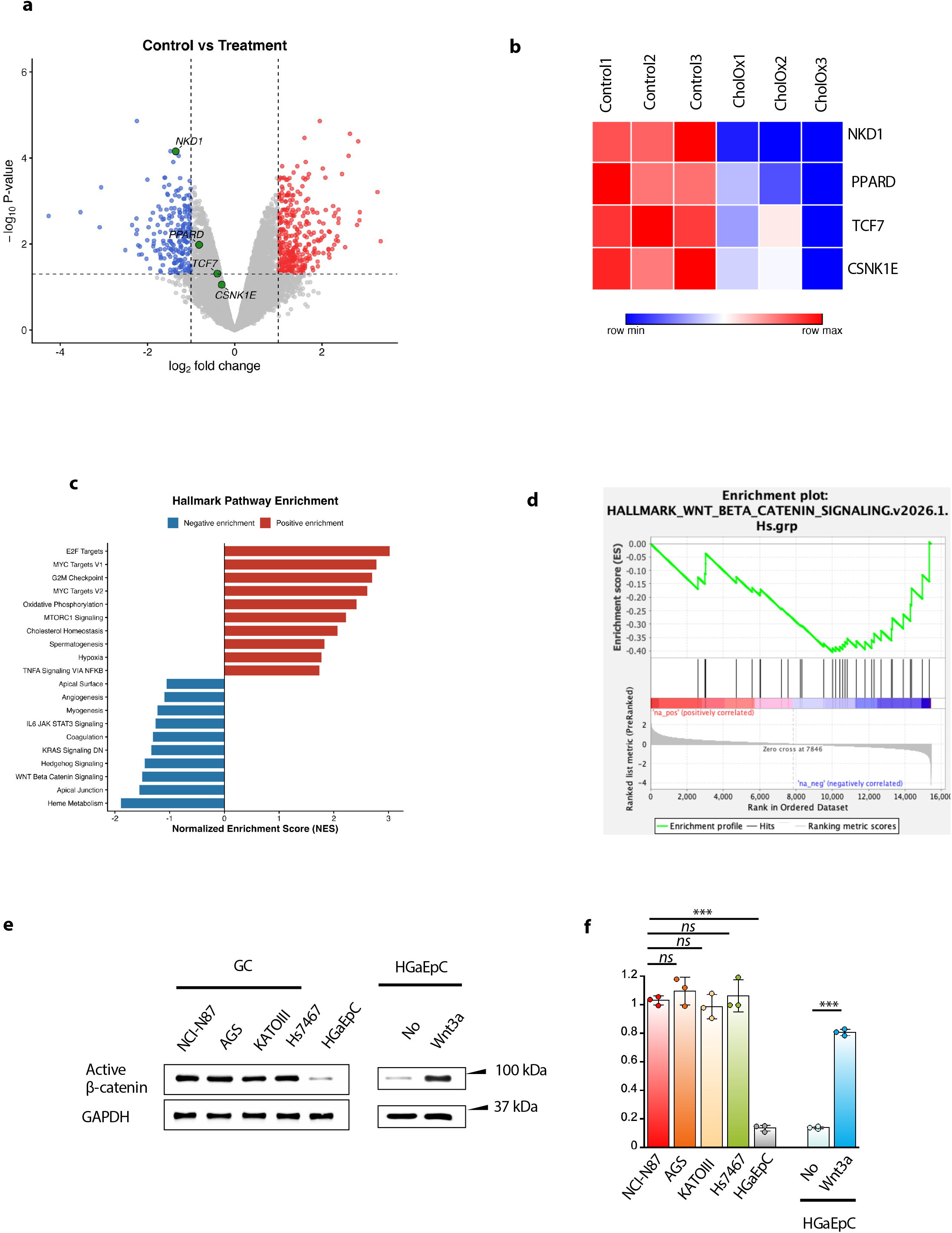
Effects of elevated IPM cholesterol on GC cell signaling activity. **(a)** Volcano plot of differentially expressed genes (DEGs) comparing control vs. cholesterol-depleted AGS cells. Highlighted genes (including *NKD1*, *PPARD*, *TCF7*, and *CSNK1E*) represent key components associated with the Wnt/β-catenin signaling cascade. **(b)** Gene Set Enrichment Analysis (GSEA) bar plot of Hallmark pathways enriched in control versus cholesterol-depleted AGS cells, displaying Normalized Enrichment Scores (NES) for both positively and negatively enriched processes. Wnt/β-catenin signaling is the most significantly disrupted signaling pathway. **(c**) GSEA enrichment plot for the Wnt/β-catenin signaling pathway (*HALLMARK WNT_BETA_CATENIN_SIGNALING*) showing a pronounced negative enrichment following cholesterol depletion. (**d**) Heatmap showing expression levels of selected Wnt/β-catenin target and pathway genes (*NKD1*, *WNT6*, *PPARD*, *TCF7*, *CSNK1E*, *AXIN1*) across replicates of control (*n* = 3) and cholesterol-depleted (CholOx) (*n* = 3) AGS cells. Relative expression is displayed as row z-scores from minimum (blue) to maximum (red). **(e)** Immunoblot analysis of active β-catenin in gastric cancer (GC) cell lines and normal gastric epithelial cells (HGaEpC). GAPDH was used as a loading control. **(f)** Quantification of active β-catenin levels, normalized to GAPDH and expressed relative to a reference cell line. All values are means ± SD. Statistical analysis was performed with unpaired Student’s t-test. *ns*, *p* > 0.05; *, *p* < 0.05; **, *p* < 0.01; ***, *p* < 0.001.

We recently reported that elevated IPM cholesterol in *Apc*-truncated CRC cells drives Wnt-independent stabilization of non-phosphorylated (active) β-catenin, leading to high constitutive β-catenin transcriptional activity (18). Notably, high baseline levels of active β-catenin were also present across all unstimulated GC cells, whereas in primary gastric epithelial cells, active β-catenin was detectable only following Wnt3a stimulation (**Fig. 3e, 3f**). Together, these findings demonstrate that elevated IPM cholesterol in GC drives constitutively active Wnt-β-catenin signaling in GC cells. Given the central role of β-catenin in the pathogenesis of gastrointestinal cancers (37–39), IPM cholesterol-mediated β-catenin signaling appears essential for GC proliferation and survival.

### Identification of ezetimibe as a site-specific modulator of inner plasma membrane cholesterol

Since elevated IPM cholesterol constitutively activates the β-catenin signaling and promotes cell proliferation in GC cells, we sought small molecule modulators that can specifically deplete IPM cholesterol as potential antitumor agents. To identify such small molecules, we performed a focused screening using representative pharmacological modulators of cholesterol import (ezetimibe), transport (OSW-1, AI-3d, U1866A), and derivatization (avasimibe) by systematically quantifying their effects on intracellular cholesterol levels in two GC cells, NCI-N87 and AGS. Inhibitors of cholesterol biosynthesis (e.g., statins) were not included because of their reported non-site-specific cytotoxicity (24).

In general, most of these inhibitors exerted relatively small effects across cell membranes (i.e., IPM, lysosomes, ER, and mitochondria) (**Fig. 4a** and **Fig. 4a**). Two exceptions were found for U1866A and ezetimibe. U1866A, an inhibitor of NPC1—which facilitates cholesterol export from lysosomes—raised the cholesterol level in the cytofacial leaflet of lysosomes by 55%, while altering cholesterol levels in all other intracellular membranes to smaller degrees, consistent with the established role of NPC1.

**Fig 4.**
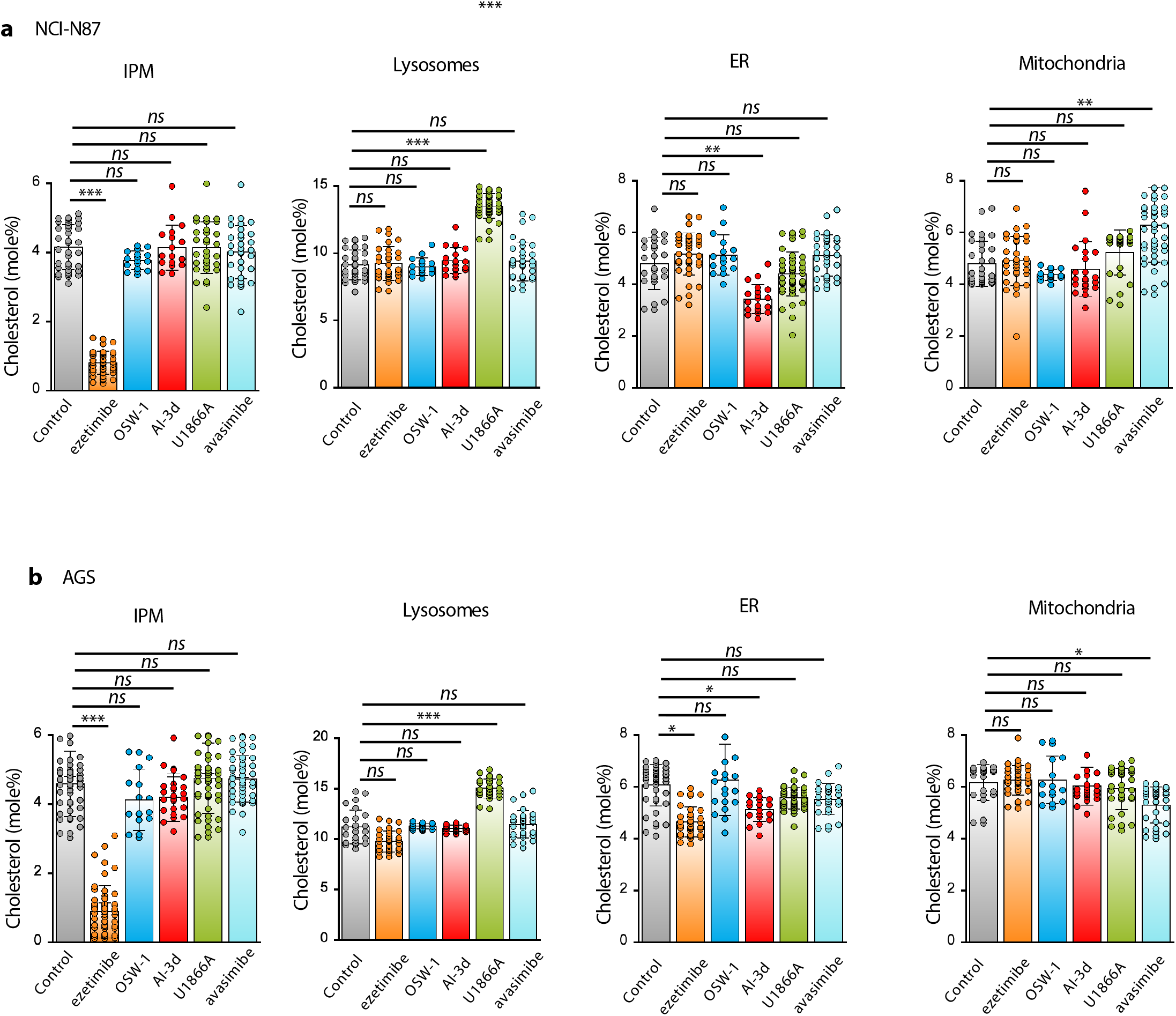
Effects of cholesterol homeostasis inhibitors on local cholesterol concentrations in NCI-N87 (a) and AGS (b) cells. Spatially resolved cholesterol quantification was performed as described in Fig. 1. Quantification was performed in triplicate (*N*=3) with different samples and *n* ≥ 6 cells per experiment. Values are means ± SD. Statistical analysis was performed with unpaired Student’s t-test. *ns*, *p* > 0.05; *, *p* < 0.05; **, *p* < 0.01; ***, *p* < 0.001.

Notably, ezetimibe uniquely and selectively reduced cholesterol levels in the IPM of the two GC cells, with minimal effects on cholesterol levels in other intracellular compartments (**Fig. 4a** and **4b**). Ezetimibe is a clinically approved inhibitor of NPC1L1, a cholesterol transporter best known for its role in intestinal cholesterol absorption. This discovery not only provides a new tool for site-specific modulation of IPM cholesterol levels in GC cells, but also yields the mechanistic insight that NPC1L1 might play a key role in regulating these levels.

### NPC1L1 serves as a cholesterol flippase in the plasma membrane to maintain high IPM cholesterol in gastric cancer cells

NPC1L1 is highly expressed in tissues connected to the digestive system, including the liver and the small intestine (35); however, recent reports have indicated that NPC1L1 is also aberrantly expressed in cancer cells and may be involved in cancer progression (36, 37). To understand how NPC1L1 is linked to cancer progression—and specifically how ezetimibe selectively reduces GC cell IPM cholesterol—we measured the effects of NPC1L1 on cholesterol distribution between the two PM leaflets of GC cells.

We first profiled NPC1L1 expression across all GC cell lines investigated. Western blot analysis revealed that GC cells exhibiting high [Chol]*_ipm_* consistently overexpressed NPC1L1 (**Fig. 5a, 5b**), corroborating previous mRNA-based bioinformatics data (38). Conversely, NPC1L1 was virtually undetectable in primary gastric epithelial cells (HGaEpC) (**Fig. 5a, 5b**), which display significantly lower [Chol]*_ipm_* (see **Fig. 1b**). To elucidate whether elevated NPC1L1 drives these uniformly high [Chol]*_ipm_* values, we performed siRNA-mediated knockdown of NPC1L1 in two representative GC cell lines, NCI-N87 and AGS. In NCI-N87 cells, knockdown markedly reduced [Chol]*_ipm_* (from 4.3 ± 0.7 to 0.6 ± 0.4 mole%) and significantly increased [Chol]*_opm_* (from 31.8 ± 0.8 to 35.2 ± 0.6 mole%), resulting in more polarized distribution of cholesterol across the PM while maintaining the same total cholesterol levels in the PM (**Fig. 5c**). The treatment did not cause statistically significant alterations of the cholesterol levels in other subcellular compartments (**Fig. 5c**). A similar trend was observed with AGS cells (**Fig. 5d**). These data suggest that NPC1L1 specifically facilitates cholesterol translocation from the OPM to the IPM. To validate this mechanism, we overexpressed NPC1L1 in non-cancerous HEK293 cells (selected due to the poor transfection efficiency of HGaEpC cells). NPC1L1 overexpression increased [Chol]*_ipm_* by 1.8-fold (i.e., from 2.2 ± 0.5 to 3.9 ± 0.5 mole%) and decreased [Chol]*_opm_* from 36.2 ± 0.5 to 33.0 ± 1.0 mole% (**Fig. 5e**), while exerting statistically insignificant alternations in other organelle membranes. The changes observed in HEK293 cells were not as dramatic as those in GC cells, presumably because the ectopically expressed NPC1L1 activity in HEK293 cells is counterbalanced by endogenous ABCA1 and ABCG1 as reported previously (16). Collectively, these results support the conclusion that NPC1L1 serves as a cholesterol flippase that shuttles cholesterol from the OPM to the IPM, and that ezetimibe specifically blocks this flippase activity.

**Fig. 5.**
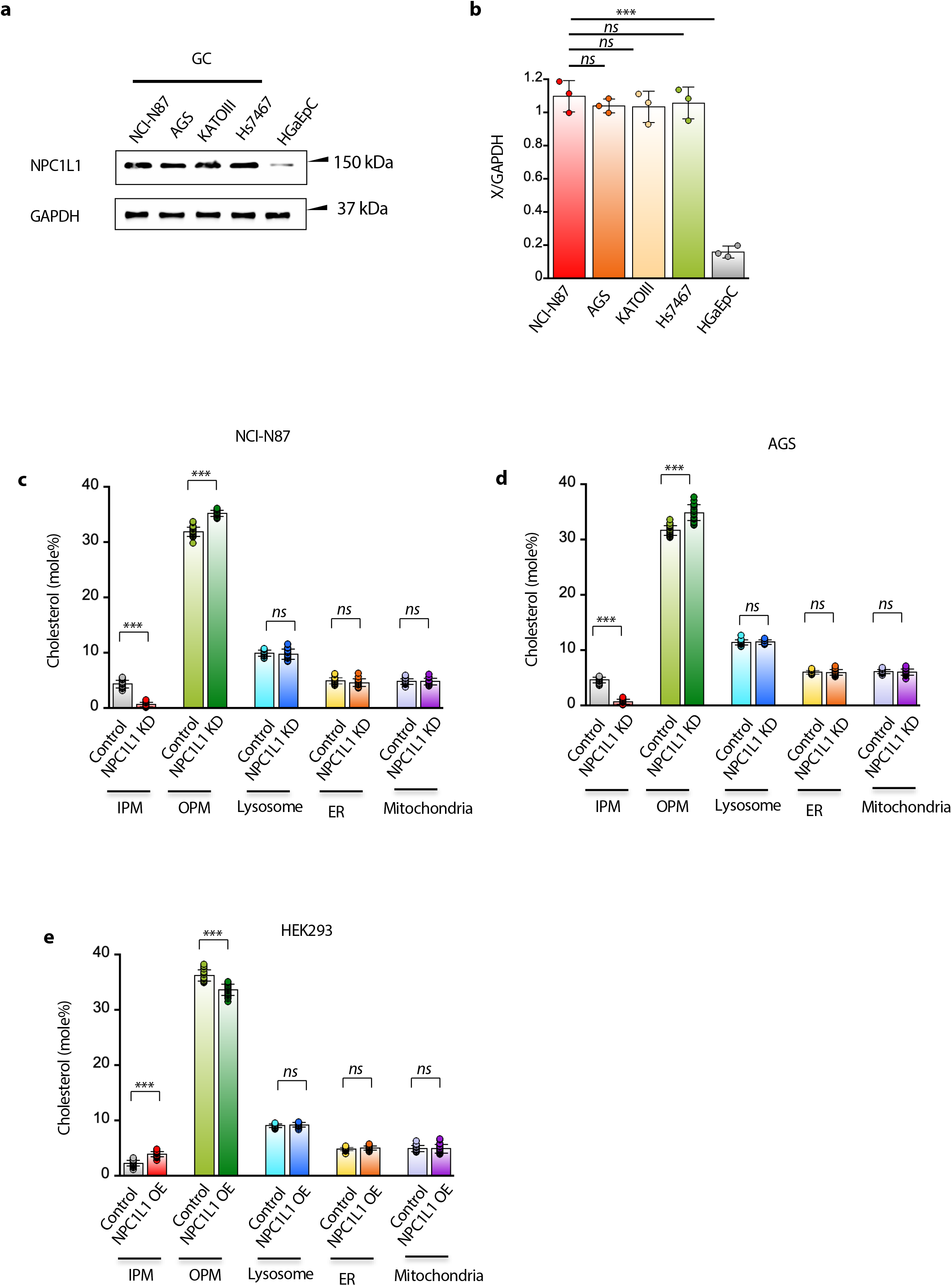
NPC1L1 controls PM cholesterol levels in GC cells. **(a)** Immunoblot analysis of NPC1L1 in gastric cancer (GC) cell lines and normal gastric epithelial cells (HGaEpC). GAPDH was used as a loading control. Representative blots from three independent experiments (n = 3) are shown. **(b)** Quantification of NPC1L1 protein levels from (a), normalized to GAPDH and expressed relative to a reference cell line. (**c-d)** Effects of siRNA-mediated knockdown (KD) of NPC1L1 on subcellular cholesterol distribution in NCI-N87 (**c**) and AGS (**d**) cells. **(e)** Effects of NPC1L1 overexpression (OE) on subcellular cholesterol distribution in HEK293 cells. For C-E, *N* = 3 and *n* = 7-10 (indicated in each plot). All values are means ± SD. Statistical analysis was performed with unpaired Student’s t-test. *ns*, *p* > 0.05; *, *p* < 0.05; **, *p* < 0.01; ***, *p* < 0.001.

### Ezetimibe suppresses IPM cholesterol-driven β-catenin signaling activity in gastric cancer cells

To determine the mechanistic link between NPC1L1 expression, IPM cholesterol levels and β-catenin signaling, we measured the effects of the siRNA-mediated knockdown of NPC1L1 on β-catenin signaling in GC cell lines. NPC1L1 knockdown, which greatly lowered [Chol]_ipm_ (see **Fig 5c, 5d**), also greatly suppressed non-phosphorylated (active) β-catenin levels in the two unstimulated GC cells, NCI-N87 and AGS (**Fig. 6a, 6b**). These results support the notion that NPC1L1-mediated upregulation of IPM cholesterol pool is responsible for sustained Wnt-independent β-catenin signaling in the GC cells. We next investigated whether ezetimibe-mediated IPM cholesterol depletion directly impacts β-catenin signaling. Treatment of four GC cell lines with ezetimibe resulted in a pronounced reduction in active β-catenin levels, as assessed by immunoblotting (**Fig. 6c**). Quantification across multiple cell lines revealed 70-80% inhibition in the active β-catenin level in the GC cells (**Fig. 6d**). Curve fitting from the dose-dependent β-catenin inhibition by ezetimibe in NCI-N87 cells yielded IC_50_ = 2.7 ± 1.3 μM (**Fig. 6e, 6f**), which is comparable to its reported IC_50_ value (3.86 μM) for the inhibition of cellular NPC1L1 activity (39). In contrast, ezetimibe had minimal effects on β-catenin levels in normal gastric epithelial cells with and without Wnt3a stimulation (**Fig. 6g, 6h**), consistent with their low NPC1L1 expression and low [Chol]_ipm_. We also determined the effects of ezetimibe on the β-catenin transcriptional output using the luciferase-based TOP-Flash assay. Ezetimibe treatment reduced β-catenin-dependent transcription in the GC cells by 60-70%, whereas a much smaller inhibitory effect was observed in the normal gastric epithelial cells (**Fig. 6i**).

**Fig. 6.**
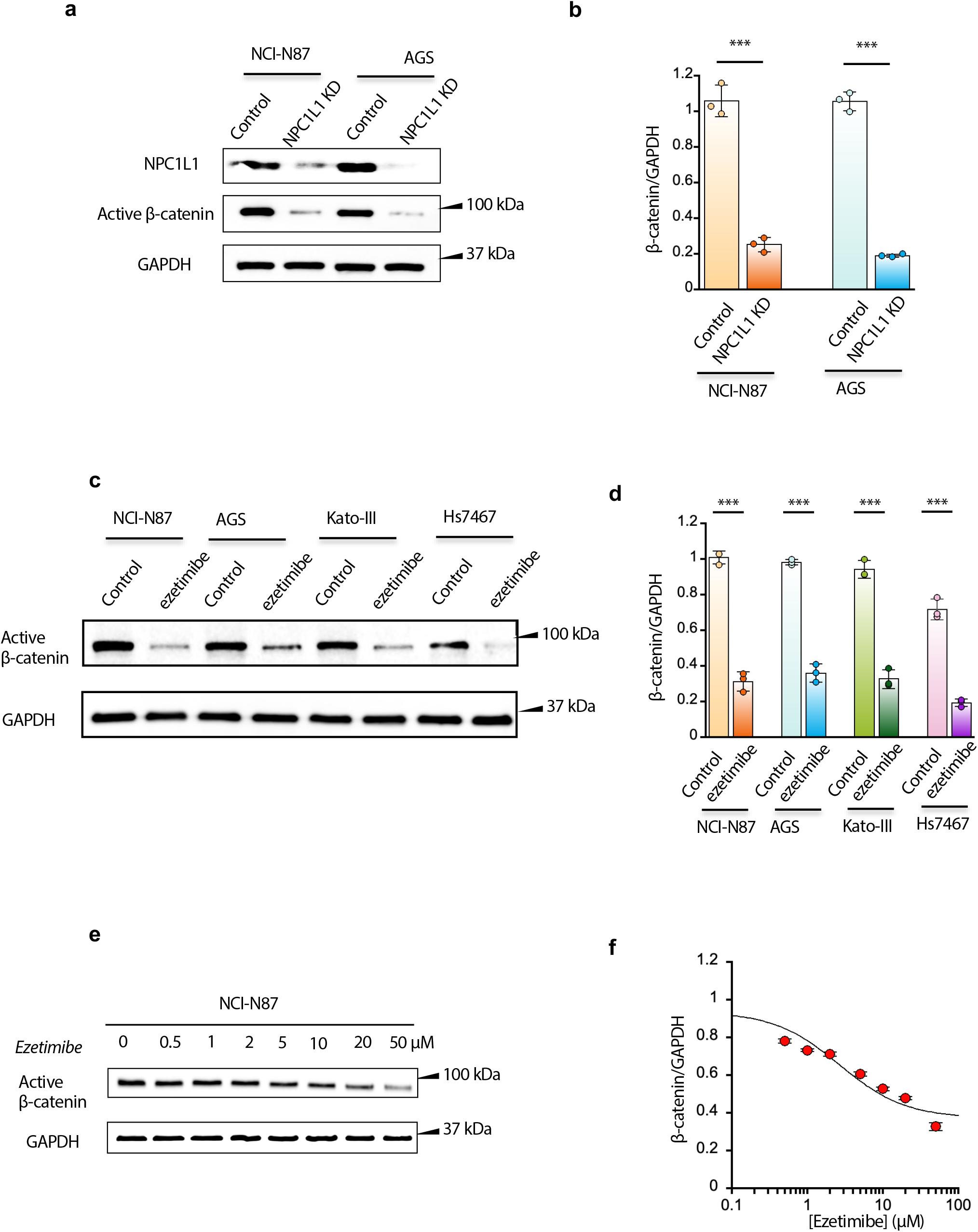
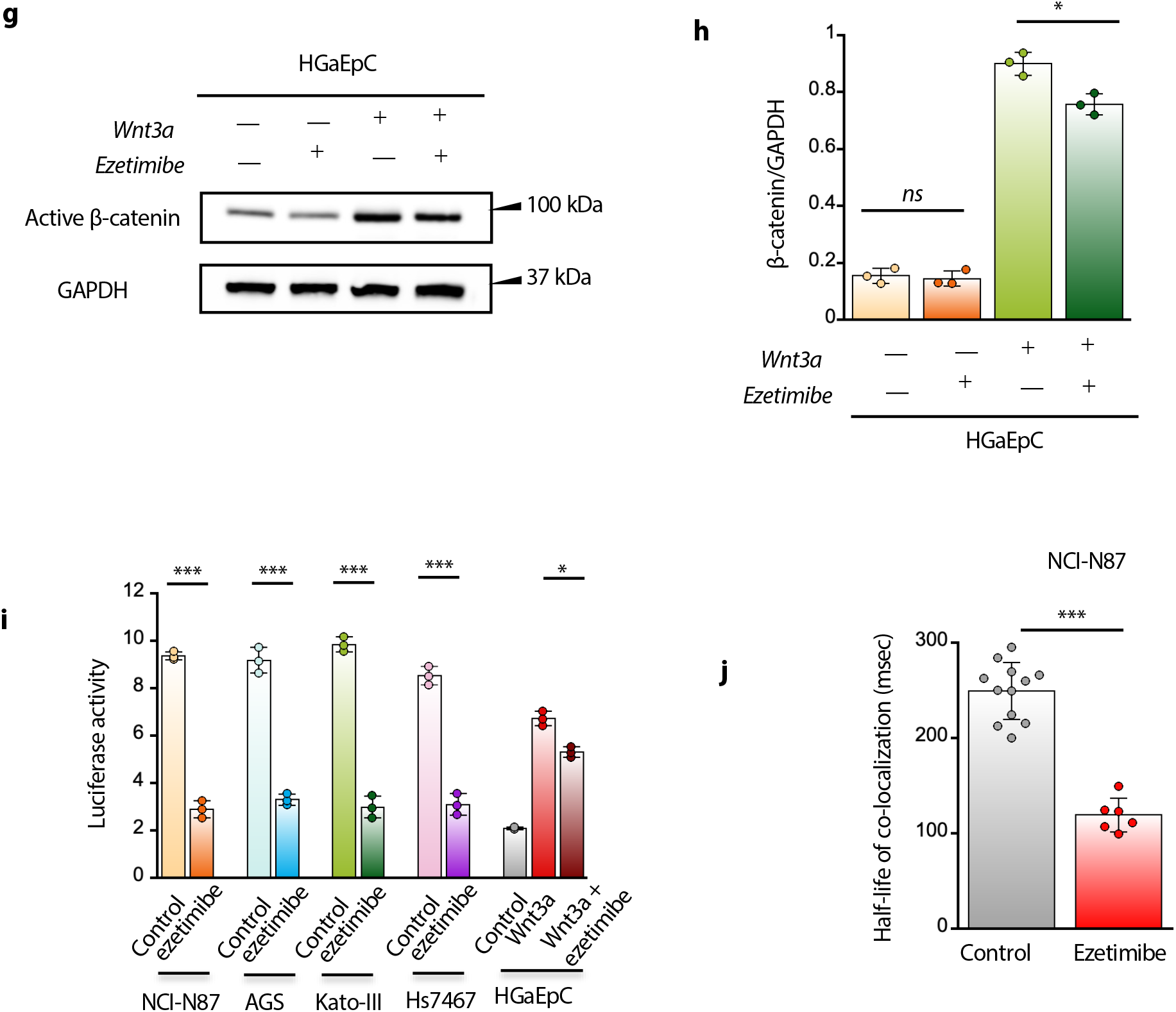
Effects of NPC1L1 downregulation on GC cell signaling activity. **(a)** Effects of siRNA-mediated NPC1L1 knockdown on the active β-catenin levels in two GC cell lines. **(b)** Quantification of **a**. **(c)** Effects of NPC1L1 inhibition by ezetimibe (20 μM, 24h) on the active β-catenin levels in four GC cell lines. **(d)** Quantification of **c**. **(e)** Dose-dependent inhibition of β-catenin by ezetimibe (0-50 μM, 24h) in NCI-N87 cells. **(f)** Quantification of **e**. The IC_50_ value was determined by non-linear least-squares analysis of the data using the equation: *y* = *y*_max_ / (1 + [inhibitor] / IC_50_) + y_min_ where *y*_max_ and *y*_min_ indicate maximal and minimal values of β-catenin/GAPDH. **(g)** Effects of NPC1L1 inhibition by ezetimibe (20 μM, 24h) on the active β-catenin levels in primary HGaEpC cells before and after Wnt3a stimulation (50 ng/ml, 12h). **(h)** Quantification of **g**. (**i**) Effects of NPC1L1 inhibition by ezetimibe (20 μM, 24h) on the β-catenin transcriptional activity (measured by the TopFlash assay) in four GC cell lines and primary HGaEpC (after 12 h stimulation with 50 ng/ml Wnt3a) cells. (**j**) Effects of NPC1L1 inhibition by ezetimibe (20 μM, 24h) on dynamic co-localization of Dvl2 and Axin 1 in NCI-N87 cells. The half-life of colocalization for the EGFP-Dvl2-Halo™-TMR-Axin1 pair was calculated from dual-color single molecule tracking analysis. **(k)** Dose-dependent inhibition of GC and primary HGaEpC cells by ezetimibe (0-50 μM, 24h). The IC_50_ values were determined by non-linear least-squares analysis of the data using the equation: *y* = *y*_max_ / (1 + [inhibitor] / IC_50_) + y_min_ where *y*_max_ and *y*_min_ indicate maximal and minimal values of relative cell viability. IC_50_ = ± (NCI-N87), (AGS), (Kato-III), (Hs7467). IC_50_ >> 30 μM for HGaEpC cells. (**l**) Inhibition of the colony formation of NCI-N87 and AGS cells by ezetimibe (20 μM, 24h). All data indicate means ± SD from three independent measurements (*N* = 3). In all immunoblot analyses, GAPDH was used as loading controls. Representative blots are shown. Active β-catenin levels were normalized to GAPDH and expressed relative to vehicle-treated controls. Statistical analysis was performed with unpaired Student’s t-test. *ns*, *p* > 0.05; *, *p* < 0.05; **, *p* < 0.01; ***, *p* < 0.001.

It was reported that elevated IPM cholesterol drives Wnt signaling in CRC cells by facilitating the formation of Wnt ligand-independent signalosomes, in which key signaling proteins, such as axin1, Dvl, FZD, and LRP5/6, dynamically interact with one another in a cholesterol-dependent manner (18). By dynamic dual-color single molecule tracking of Dvl and axin1, we monitored constitutive Wnt-independent formation of Wnt signalosome in NCI-N87 cells. Even without Wnt3a stimulation, Dvl and axin1 exhibited a significant degree of dynamic co-localization in NCI-N87 cells, as reported in CRC cells (18). Notably, this Wnt-independent co-localization of Dvl and axin1 was suppressed by ezetimibe to the basal level, indicating destabilization of Wnt receptor/signalosome assemblies at the PM by ezetimibe (**Fig. 6j**). Together, these data indicate that NPC1L1-mediated IPM cholesterol elevation in GC cells drives Wnt-β-catenin signaling by facilitating the formation of ligand-independent Wnt signaling complexes, which is blocked by ezetimibe.

### Ezetimibe potently and specifically inhibits gastric cancer cell viability

We next examined whether suppression of β-catenin signaling translated into reduced cancer cell viability in both two-dimensional (2D) and three-dimensional (3D) cell culture models. Treatment with ezetimibe significantly decreased the viability of all four GC cell lines in a dose-dependent manner (**Fig. 7a**). Notably, this phenotypic response closely mirrored the concentration thresholds required for β-catenin inhibition (**Fig. 6e, 6f**), suggesting a direct functional link. In contrast, primary gastric epithelial cells remained largely resilient to ezetimibe at concentrations that potently suppressed cancer cell growth (**Fig. 7a**). This selective cytotoxicity aligns with their low baseline NPC1L1 expression (**Fig. 5a**) and correspondingly diminished IPM cholesterol level (**Fig. 1b**), highlighting a favorable therapeutic window. To extend these findings to a more physiologically relevant architecture, we evaluated the effects of ezetimibe on 3D GC cultures; the drug robustly suppressed the anchorage-independent growth of NCI-N87 and AGS cells on agar (**Fig. 7b**).

**Fig. 7.**
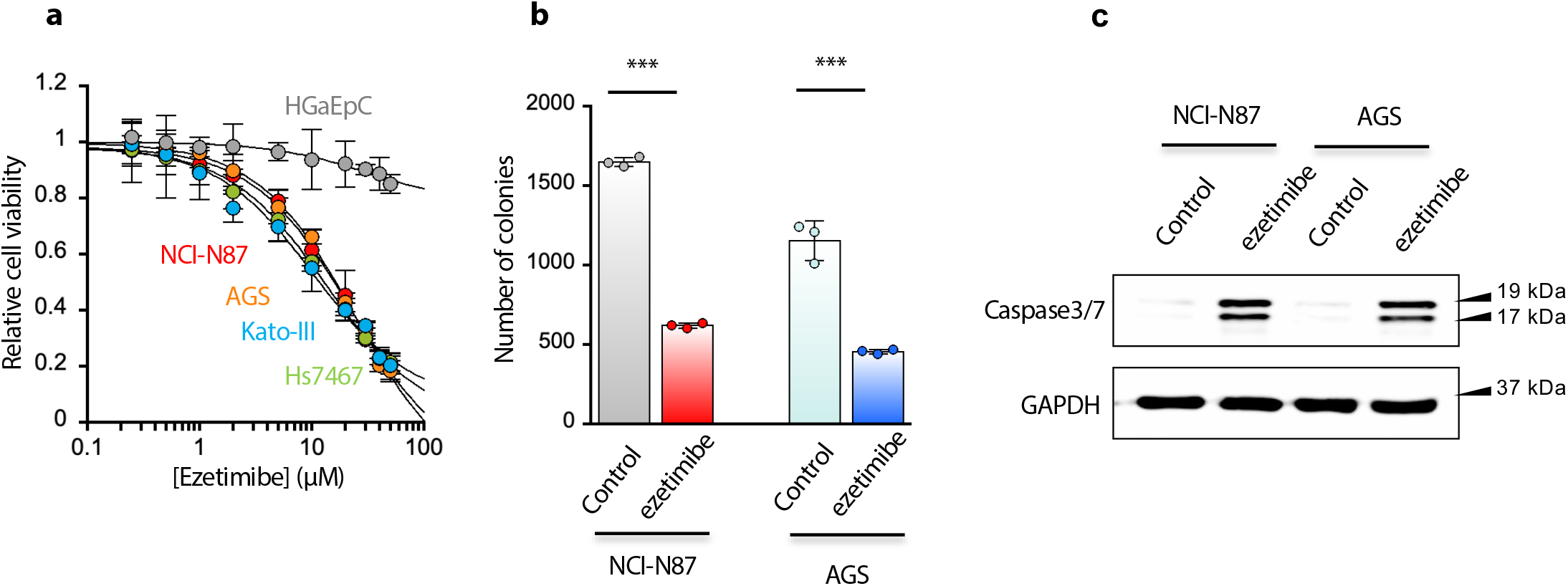
Effects of ezetimibe on GC cell viability. **(a)** Dose-dependent inhibition of GC and primary HGaEpC cells by ezetimibe (0-50 μM, 24h). The IC_50_ values were determined by non-linear least-squares analysis of the data using the equation: *y* = *y*_max_ / (1 + [inhibitor] / IC_50_) + y_min_ where *y*_max_ and *y*_min_ indicate maximal and minimal values of relative cell viability. IC_50_ = 10.4 ± 2.2 (NCI-N87), 13.4 ± 1.3 (AGS), 22.1 ± 2.5 (Kato-III), 24.0 ± 3.0 (Hs7467) μM. IC_50_ >> 30 μM for HGaEpC cells. (**b**) Inhibition of the colony formation of NCI-N87 and AGS cells by ezetimibe (20 μM, 24h). (**c**) Induction of apoptosis in NCI-N87 cells by ezetimibe (20 μM, 24h) as monitored by caspase 3 cleavage. All data indicate means ± SD from three independent measurements (*N* = 3). In all immunoblot analyses, GAPDH was used as loading controls. Representative blots are shown. Active β-catenin levels were normalized to GAPDH and expressed relative to vehicle-treated controls. Statistical analysis was performed with unpaired Student’s t-test. *ns*, *p* > 0.05; *, *p* < 0.05; **, *p* < 0.01; ***, *p* < 0.001.

To determine whether this loss of viability was driven by apoptotic cell death, we assessed caspase activation cascade dynamics following ezetimibe exposure. Immunoblot analysis revealed a robust increase in cleaved caspase-3 in GC cells (**Fig. 7c**). Taken together, these data indicate that ezetimibe suppresses tumor cell growth through the coordinated downregulation of Wnt/β-catenin signaling and the active induction of apoptosis, rather than via non-specific, off-target toxicity.

### High expression of NPC1L1 in gastric cancer is correlated with poor prognosis

To explore whether NPC1L1 might contribute to cholesterol-dependent signaling in cancer cells, we performed a comprehensive bioinformatic analysis of NPC1L1 expression profiles across a diverse panel of human malignancies. Based on the newly identified function of NPC1L1 as a cholesterol flippase that controls IPM cholesterol levels and cell proliferation rates, we hypothesized that its dysregulation might trigger oncogenic pathways.

Consistent with recent independent bioinformatics datasets (38), our pan-cancer screening revealed that *NPC1L1* expression in primary tumors is heterogeneous but peaks prominently in gastrointestinal malignancies, showing the highest transcript levels in in GC and pancreatic ductal adenocarcinoma (**Fig. 8a**). This specific enrichment suggests that tumors arising from tissues structurally involved in nutrient absorption and cholesterol homeostasis may hijack NPC1L1 to meet their heightened metabolic and proliferative demands. A detailed comparison of NPC1L1 mRNA expression confirmed a significant enrichment in primary gastric tumor tissues compared to adjacent normal tissues (**Fig. 8b**).

**Fig. 8.**
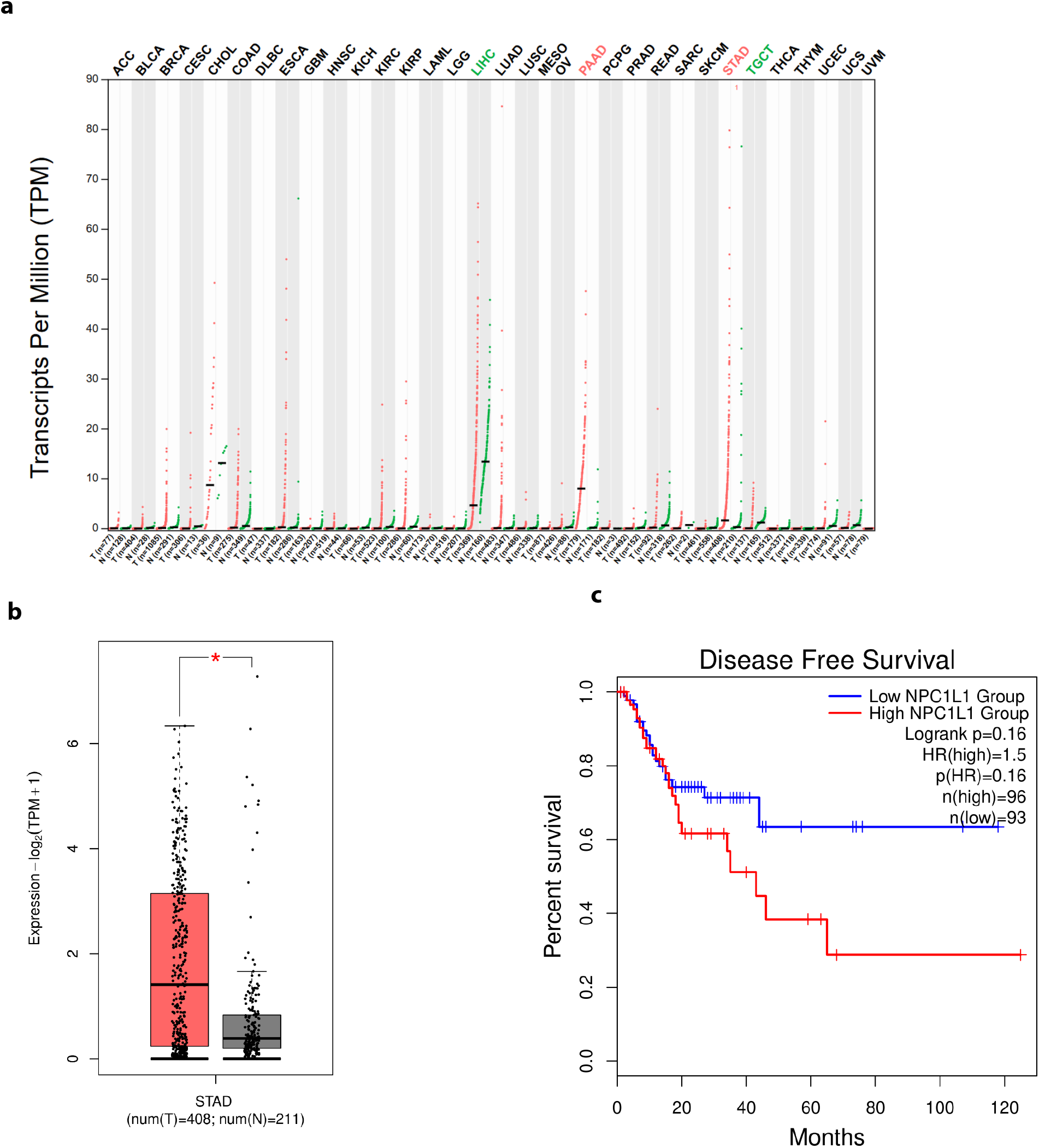
Pan-cancer expression profile and clinical prognostic value of NPC1L1. **(a)** Pan-Cancer Expression Profile of NPC1L1: Comparative analysis of relative NPC1L1 mRNA expression levels between tumor (T; red) and adjacent normal (N; green) tissues across multiple malignancies using integrated data from the TCGA and GTEx databases. Expression values are quantified as Transcripts Per Million (TPM). Individual data points represent independent patient samples, with sample sizes (n) specified for each cohort along the horizontal axis. Cancer types exhibiting noteworthy expression alterations are highlighted in color-coded text (e.g., green for LIHC and TGCT; red for PAAD and STAD). **(b)** Differential NPC1L1 Expression in Stomach Adenocarcinoma (STAD): Detailed comparison of NPC1L1 mRNA expression between STAD tumor tissues (*n* =408, red box) and normal gastric tissues (*n* = 211, grey box). To normalize the distribution and account for variance, expression levels are plotted on a log-transformed scale: log2 (TPM+1). Horizontal center lines within the box plots represent the median expression value, while individual dots represent individual tissue samples. *, p<0.05. **(c)** Kaplan-Meier Disease-Free Survival Analysis: Kaplan-Meier survival curves estimating the impact of NPC1L1 expression on Disease-Free Survival over a follow-up period of up to 120 months in GC patients from the TCGA database. Patients were stratified into a High NPC1L1 Expression group (*n* = 96, red curve) and a Low NPC1L1 Expression group (*n* = 93, blue curve). Small vertical tick marks along the curves denote censored observations. Statistical differences between the cohorts were evaluated using the log-rank test (*p* = 0.16), with the calculated Hazard Ratio (HR=1.5) indicating a trend toward shortened survival in the high-expression subgroup. **Abbreviations: ACC:** Adrenocortical carcinoma; **BLCA:** Bladder Urothelial Carcinoma; **BRCA:** Breast invasive carcinoma; **CESC:** Cervical squamous cell carcinoma and endocervical adenocarcinoma; **CHOL:** Cholangiocarcinoma; **COAD:** Colon adenocarcinoma; **DLBC:** Diffuse Large B-cell Lymphoma; **ESCA:** Esophageal carcinoma; **GBM:** Glioblastoma multiforme; **HNSC:** Head and Neck squamous cell carcinoma; **KICH:** Kidney Chromophobe; **KIRC:** Kidney renal clear cell carcinoma; **KIRP:** Kidney renal papillary cell carcinoma; **LAML:** Acute Myeloid Leukemia; **LGG:** Brain Lower Grade Glioma; **LIHC:** Liver hepatocellular carcinoma; **LUAD:** Lung adenocarcinoma; **LUSC:** Lung squamous cell carcinoma; **MESO:** Mesothelioma; **OV:** Ovarian serous cystadenocarcinoma; **PAAD:** Pancreatic adenocarcinoma; **PCPG:** Pheochromocytoma and Paraganglioma; **PRAD:** Prostate adenocarcinoma; **READ:** Rectum adenocarcinoma; **SARC:** Sarcoma; **SKCM:** Skin Cutaneous Melanoma; **STAD:** Stomach adenocarcinoma; **TGCT:** Testicular Germ Cell Tumors; **THCA:** Thyroid carcinoma; **THYM:** Thymoma; **UCEC:** Uterine Corpus Endometrial Carcinoma; **UCS:** Uterine Carcinosarcoma; **UVM:** Uveal Melanoma.

Finally, to determine whether this expression pattern translates to clinical outcomes, we conducted a survival analysis across patient cohorts. Elevated *NPC1L1* expression showed a clear trend toward shortened disease-free survival in GC patients, although this did not reach statistical significance (**Fig. 8c**; *p = 0.16, HR =1.5*), likely due to cohort size limitations. Nevertheless, these collective findings position NPC1L1 as a promising candidate biomarker in GC and provide a compelling rationale for targeting it to disrupt cholesterol-dependent pro-tumorigenic signaling and mitigate tumor progression.

## DISCUSSION

NPC1L1 is highly expressed in tissues connected to the digestive system, including the liver and the small intestine (35). While the role of NPC1L1 in intestinal cholesterol absorption is well established, its precise mechanism of action remains elusive. It was initially reported that NPC1L1 facilitates the entry of cholesterol into enterocytes primarily through clathrin-mediated endocytosis and that ezetimibe blocks this cholesterol-induced internalization of NPC1L1 (40). This notion was later challenged by a report indicating that cholesterol uptake by NPC1L1 does not require endocytosis and that ezetimibe interferes with NPC1L1’s cholesterol absorption activity without blocking NPC1L1 internalization (41). Consistently, three recent structural studies (39, 42, 43) demonstrated that NPC1L1 contains an intramolecular hydrophobic tunnel that can serve as a conduit for transbilayer cholesterol transport. Our study provides definitive evidence that NPC1L1 can indeed function as a cholesterol flippase, actively shuttling cholesterol from the outer to the inner leaflet of the plasma membrane.

Recent reports have indicated that NPC1L1 is also aberrantly expressed in cancer (36, 37). Although growing evidence indicates that NPC1L1 plays a significant role in the development, progression, and therapeutic resistance of several malignancies (36, 37)—including GC, CRC, pancreatic cancer, prostate cancer, and breast cancer—the exact molecular mechanisms underlying its functional dynamics in tumor biology remain poorly understood. Multiple bioinformatics analysis, including ours, show that NPC1L1 is most abundantly expressed in GC and pancreatic cancer. For instance, a recent study showed that NPC1L1 in pancreatic cancer cells not only facilitates tumor proliferation by promoting cholesterol import, but also serves as an immune checkpoint to suppress CD8+ T cell activation (44). Crucially, our bioinformatics analysis reveals that NPC1L1 is enriched in primary gastric tumors compared with neighboring tissues and exhibits a strong trend toward an inverse correlation with patient survival.

Our biophysical and cell biology studies provide mechanistic insight into how NPC1L1 drives GC cell proliferation and survival. Cholesterol has long been known as an essential structural component of cellular membranes, but growing evidence indicates that cholesterol also functions as a site-specific signaling lipid. In cancer cells where lipid metabolism is frequently reprogrammed, the altered distribution of cholesterol and its derivatives across cellular compartments has been liked to malignancies (5). In this study, we demonstrate that NPC1L1 facilitates elevation of IPM cholesterol in GC cells. This cholesterol elevation drives stimulus-independent, cell-intrinsic, constitutive activation of β-catenin signaling, ultimately leading to amplified cell proliferation. We recently reported that elevated IPM cholesterol in CRC cells constitutively activates β-catenin signaling by inducing PM recruitment of a scaffolding protein Dvl that coordinates Wnt ligand-independent formation of β-catenin signalosomes (18). Our biochemical and single molecule imaging studies indicate that this identical mechanism operates in GC cells.

We previously reported that ABCA1 (and ABCG1) controls IPM cholesterol levels by serving as an ATP-dependent lipid translocase (floppase) that shuttles cholesterol from the IPM to the OPM (16). Furthermore, the expression level of ABCA1 is inversely correlated with IPM cholesterol levels in many cancer cells (18). Here, the direct correlation observed between NPC1L1 expression levels and IPM cholesterol levels in GC and other cells strongly supports the conclusion that NPC1L1 facilitates the reverse translocation of cholesterol from the OPM to the IPM. This model aligns with recent structural studies of NPC1L1 (39, 42, 43). Notably, a cholesterol flippase that specifically facilitates such inward translocation has not been previously reported.

Previous efforts to target cholesterol metabolism in cancer have largely focused on inhibiting *de novo* cholesterol biosynthesis, most notably through the use of statins (9). While statins effectively reduce the total cellular cholesterol level, their effects on oncogenic signaling pathways have been inconsistent, and the clinical benefit in solid tumors remains unestablished (10). Presumably, changes in total cholesterol mass do not directly reflect the availability of cholesterol within specific membrane compartments that directly regulate cancer signaling. The fact that IPM cholesterol serves as a key site-specific pro-tumorigenic signal in multiple cancers establishes it as an attractive target for site-specific cholesterol depletion cancer therapy. Our systematic analysis of subcellular cholesterol pools shows that unlike other pharmacological inhibitors of cholesterol transport and derivatization that have relatively uniform effects on subcellular cholesterol distribution, ezetimibe selectively and effectively depletes IPM cholesterol by inhibiting flippase function of NPC1L1. As such, this represents the first example of successful site-specific depletion of IPM cholesterol in cancer cells. To date, specific pharmacological modulators of ABCA1 and ABCG1 floppase activity have not been reported.

Ezetimibe is an FDA-approved lipid-lowering agent widely prescribed to reduce low-density lipoprotein cholesterol and mitigate cardiovascular risk; however, its clinical utility has not yet been extended to oncology. In *in vitro* settings and preclinical animal models, ezetimibe has demonstrated robust antineoplastic efficacy, suppressing the progression of various malignancies—including breast, liver, and prostate cancers—primarily by blocking angiogenesis or inducing apoptosis (45, 46). Despite these promising laboratory findings, ezetimibe has proven largely ineffective as a standalone cancer therapy in clinical settings. This stark translational discrepancy likely stems from a critical mismatch in pharmacokinetics and experimental design. Pharmacologically, ezetimibe is designed to act locally within the brush border membrane of the small intestine, where it selectively inhibits the NPC1L1 transporter to block dietary and biliary cholesterol absorption. Consequently, while the drug achieves high, effective concentrations within the gastrointestinal tract, its systemic bioavailability in the bloodstream is low (47). Furthermore, many preclinical studies establishing ezetimibe’s anti-tumor efficacy were conducted using rodent models maintained on high-fat, high-cholesterol diets. In these specific hyperlipidemic models, ezetimibe exerts a dramatic, systemic indirect effect: by shutting down intestinal absorption, it profoundly depletes circulating cholesterol levels, thereby starving peripheral tumors of the exogenous lipids they rely on to fuel rapid proliferation (37). However, human cancer patients typically maintain normal or stable circulating lipid profiles. In a clinical scenario, a standard oral dose of ezetimibe does not lower systemic cholesterol aggressively enough to induce metabolic starvation in peripheral tumors, nor does it achieve a high enough serum concentration to directly bind and inhibit the low levels of NPC1L1 expressed ectopically on tumor cells outside the gut (37). Due to these pharmacokinetic complexities, *in vivo* testing of ezetimibe using GC animal models was not included in this study. Instead, the current work focuses on biophysical and mechanistic investigation of NLC1L1, which can provide a blueprint for future drug development tailored for GC.

Several important questions remain. The precise molecular mechanisms by which NPC1L1 regulates IPM cholesterol distribution in cancer cells are not yet fully understood, nor is it known whether other oncogenic signaling pathways exhibit similar dependencies on spatially restricted cholesterol pools. Additionally, while our data supports a shared cholesterol-dependent mechanism in GC cell lines, future studies will be required to define tumor-specific adaptations within this broader framework. Exploring combination strategies in which site-specific cholesterol modulation is paired with other targeted or immune-based therapies may further enhance therapeutic efficacy.

In summary, this study identifies IPM cholesterol as a critical and druggable regulator of Wnt– β-catenin signaling in GC. By repurposing ezetimibe to selectively target this cholesterol pool, we establish a paradigm in which site-specific lipid modulation destabilizes oncogenic signaling complexes, suppresses tumor growth, and spares normal tissues. These findings highlight the broader importance of membrane organization in cancer cell signaling and suggest that targeting lipid-dependent signaling dynamics represents a highly promising avenue for future oncological therapeutics.

## DATA AVAILABILITY STATEMENT

All data are contained within the manuscript. The raw data will be shared upon request: Contact Wonhwa Cho (University of Illinois Chicago).

## CONFLICT OF INTEREST STATEMENT

The authors declare no conflict of interest with the contents of the articles.

## ACKNOWLEDGEMENTS

This work was supported by a grant from the National Institutes of Health (R35GM122530).

## Notes

### Competing Interest Statement

The authors have declared no competing interest.

